# Zika Virus in Salivary Glands of Five Different Species of Wild-Caught Mosquitoes from Mexico

**DOI:** 10.1101/151951

**Authors:** Darwin Elizondo-Quiroga, Aarón. Medina-Sánchez, Jorge M. Sánchez-González, Kristen Allison Eckert, Erendira Villalobos-Sánchez, Antonio Rigoberto Navarro-Zúñiga, Gustavo Sánchez-Tejeda, Fabián Correa-Morales, Cassandra González-Acosta, Armando E. Elizondo-Quiroga, Mexican Network for Virology, Secretaría de Salud of Mexico

## Abstract

Zika virus (ZIKV) is a mosquito-borne virus and *Aedes agypti* has been mentioned as the main vector of the disease. Other mosquito species in the *Aedes* and *Culex* genera have been suggested to have the potential for being competent vectors, based on experimental exposition of mosquitoes to an infectious blood meal containing ZIKV. Here, we report the isolation in cell culture of ZIKV from different body parts of wild-caught female mosquitoes (*Ae. aegypti*, *Ae. vexans, Culex quinquefasciatus*, *Cx. coronator*, and *Cx. tarsalis*) and whole male mosquitoes (*Ae. aegypti* and *Cx. quinquefasciatus*) in Mexico. Importantly, the virus was isolated from the salivary glands of all of these mosquitoes, strongly suggesting that these species are potential vectors for ZIKV.

## Introduction

Brazil was the first country in the Western Hemisphere to report Zika virus (ZIKV) in 2015^1^, but the transmission has spread to more than 50 countries and territories in the region^2^. As of November 2016, there were 6,474 confirmed cases of ZIKV infection, including 3,663 pregnant women, in 21 states of Mexico^3^.

ZIKV is a member of the *Flaviviridae* family and the genus *Flavivirus*; the presumptive main vector of the virus is *Aedes aegypti* (L.), and laboratory studies have demonstrated its ability to acquire and potentially transmit the virus in experimentally fed mosquitoes with infected blood^4,5^. However, it has been recently shown that other mosquito species may also transmit the virus in laboratory conditions, including *Ae. vexans* (Meigen)^6, 7^, and mosquitoes in the *Culex* genus, such as *Culex quiquefasciatus* Say ^8,9^.

On the other hand, it has been reported that *Ae. aegypti* and *Ae. albopictus* (Skuse) are not competent vectors to transmit ZIKV, although they are susceptible to infection^10,11^. To understand the vector competence of different mosquito species in the metropolitan area of Guadalajara, in the State of Jalisco, Mexico, we collected mosquitoes inside houses, in neighborhoods where at least one confirmed or probable case of ZIKV in humans, had been reported by the health authorities of Mexico.

Personnel of the Entomological Research Unit of the Jalisco State Public Health Department collected mosquitoes over 5 days, from September to November 2016 in 3 different municipalities (18 blocks in 4 neighborhoods) of the metropolitan area of Guadalajara (Figure 1). Five-hundred and seventy-nine mosquitoes, representing 2 genera (*Aedes* and *Culex*) and 6 species (*Ae. aegypti, Ae. epactius* Dyar and Knab*, Ae. vexans, Cx. quinquefasciatus, Cx. coronator* Dyar and Knab, and *Cx. tarsalis* Coquillett, were collected; the mosquitoes were separated in the laboratory by species and sex into pools of maximum 25 insects each. All 579 mosquitoes in 179 pools (Table 1; Figure 1) were processed for virus isolation at CIATEJ (Centro de Investigación y Asistencia en Tecnologia y Diseño del Estado de Jalisco A.C.).

**Figure 1.**
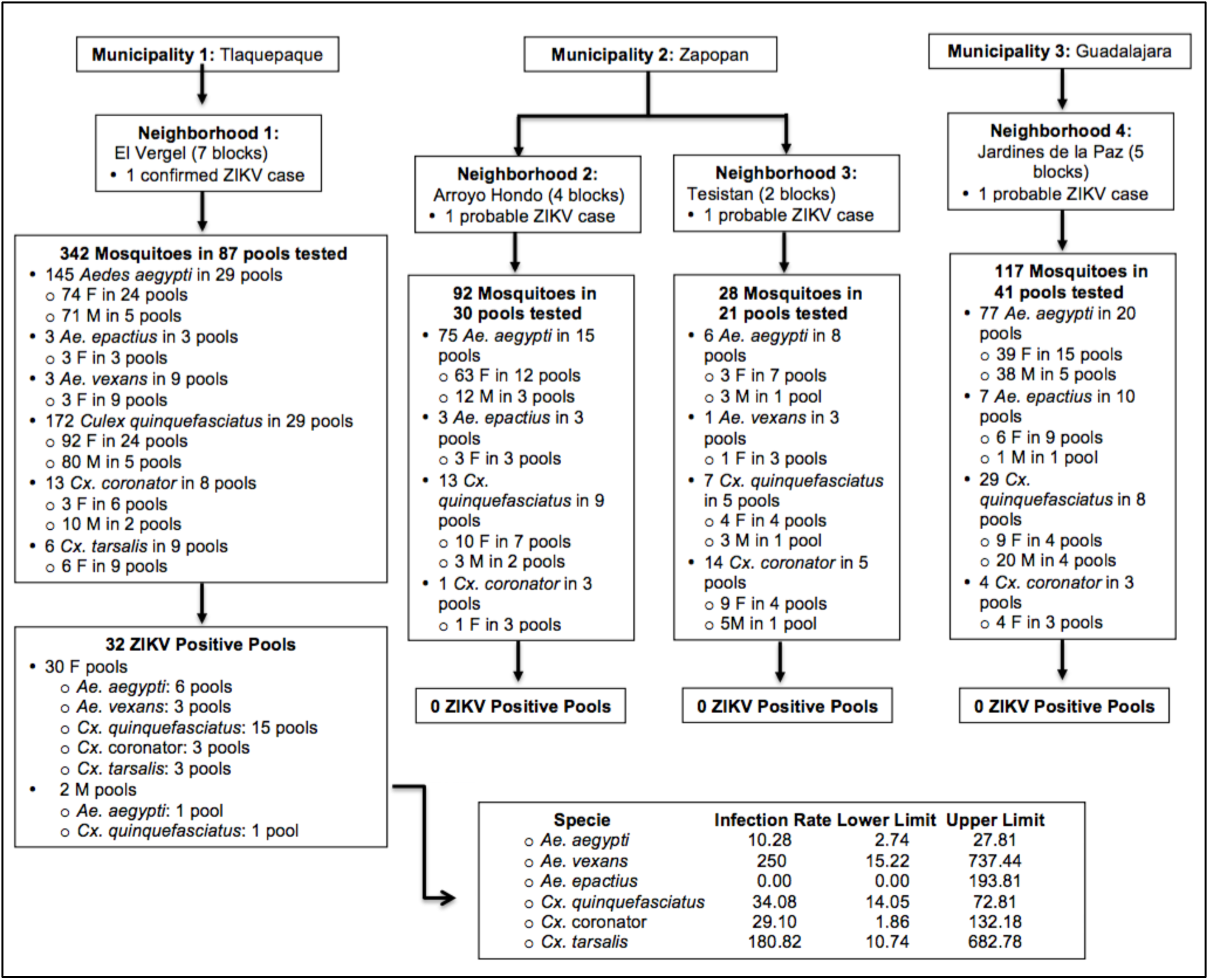
Flow chart of mosquito collection, findings and infection rates in the metropolitan area of Guadalajara, Jalisco, Mexico. F = Female; M = Male; ZIKV = Zika Virus

**Table 1.** Summary of mosquito species collected in neighborhoods from the metropolitan area of Guadalajara, Jalisco, Mexico.

|  | Female Mosquitoes |  |  | Male Mosquitoes |  |
| --- | --- | --- | --- | --- | --- |
| Genus and Species | No. mosquitoes | No. pools collected | No. pools of body parts | No. mosquitoes | No. pools collected |
| *Aedes aegypti* (n = 303) | 179 | 20 | 58 | 124 | 14 |
| *Aedes epactius* (n = 13) | 12 | 6 | 15 | 1 | 1 |
| *Aedes vexans* (n = 4) | 4 | 4 | 12 | 0 | 0 |
| *Culex quinquefasciatus* (n = 221) | 115 | 17 | 39 | 106 | 12 |
| *Culex coronator* (n = 32) | 17 | 6 | 16 | 15 | 3 |
| *Culex tarsalis* (n = 6) | 6 | 3 | 9 | 0 | 0 |
| **Total (n = 579)** | **333** | **56** | **149** | **246** | **30** |

Thirty pools of mosquitoes’ body parts (salivary glands, midguts, and rest of the bodies) from 10 pools of the dissected female mosquitoes (*Ae. aegypti*: 2 pools, *Ae. vexans*: 1 pool, *Cx. quinquefasciatus*: 5 pools, *Cx. coronator*: 1 pool, and *Cx. tarsalis*: 1 pool) and 2 pools of male mosquitoes (*Ae. aegypti* and *Cx. quinquefasciatus*), representing 5 of the 6 species collected for both genera, yielded virus isolates, with a CPE observed between 1-5 days post inoculation (dpi) in C6/36 cells (Table 2). All positive pools were from the Vergel neighborhood of Tlaquepaque (Figure 1). The isolates obtained in C6/36 cells were confirmed to infect Vero cells, and all isolates were identified as ZIKV by RT-qPCR; CT’s values for all infected cultures ranged between 13 and 16 cycles. Importantly, the female mosquitoes’ salivary glands of three *Culex species: Cx. coronator*, *Cx. Tarsalis,* and *Cx. quinquefasciatus* and two *Aedes* species: *Ae. vexans* and *Ae. aegypti* were found to be positive. This is the first report, as far as we know, that shows the presence of ZIKV in the salivary glands of wild-caught female mosquitoes from these species.

**Table 2.** Zika virus positive mosquito species and time of cytopathic effect appearance.

| **Mosquitoes Genus**<br>**and Species** | Number of<br>mosquitoes | Pools positive<br>for ZIKV<br>isolation | CPE (Dpi) |
| --- | --- | --- | --- |
| *Aedes aegypti* | 25 | Male | 2 |
| *Ae. aegypti* | 8 | SG | 3 |
|  |  | MG | 3 |
|  |  | B | 3 |
| *Ae. aegypti* | 3 | SG | 3 |
|  |  | MG | 3 |
|  |  | B | 3 |
| *Ae. vexans* | 1 | SG | 3 |
|  |  | MG | 3 |
|  |  | B | 3 |
| *Culex quinquefasciatus* | 25 | Male | 2 |
| *Cx. quinquefasciatus* | 16 | SG | 3 |
|  |  | MG | 3 |
|  |  | B | 3 |
| *Cx. quinquefasciatus* | 20 | SG | 3 |
|  |  | MG | 5 |
|  |  | B | 3 |
| *Cx. quinquefasciatus* | 25 | SG | 3 |
|  |  | MG | 3 |
|  |  | B | 3 |
| *Cx. quinquefasciatus* | 9 | SG | 1 |
|  |  | MG | 1 |
|  |  | B | 2 |
| *Cx. quinquefasciatus* | 3 | SG | 1 |
|  |  | MG | 3 |
|  |  | B | 2 |
| *Cx. coronator* | 2 | SG | 3 |
|  |  | MG | 3 |
|  |  | B | 3 |
| *Cx. tarsalis* | 3 | SG | 4 |
|  |  | MG | 4 |
|  |  | B | 1 |

SG = Salivary Gland; MG = Midgut; B Body; CPE=Cytopathic effect; Dpi=
Days post-inoculation

Of interest, we found the highest infection rate (IR) in *Ae. vexans* (250) and *Cx. tarsalis* (180.82) and the lowest in *Ae. aegypti* (10.28) (Figure 1), suggesting that the latter specie could not be the best vector for ZIKV, at least in the State of Jalisco, Mexico. These results are in accordance with previous publications where the vector competence of *Ae. aegypti* was experimentally evaluated using mosquitoes from other regions of the Americas^10^, as well as from Senegal^11^.

Most pools analyzed showed a CPE after 3 dpi, but 2 pools of salivary glands of *Cx. quinquefasciatus* showed a CPE 1 dpi (Table 2). These findings support previously published results that suggest that the *Cx quinquefasciatus* mosquito is a potential vector to transmit ZIKV^8,9^. On the other hand, the results presented in this work are discordant with previous publications reporting *Culex* spp. as bad ZIKV vectors; for instance, a report using North American mosquito colonies maintained by decades in the laboratory^13^, and in mosquitoes from Rio de Janeiro Brazil, where it was found that *Cx. quinquefasciatus* is not competent to transmit the local strain of ZIKV^14^. These observations could be explained by the genetic variability of the mosquito populations, as previously suggested^11,15^. Hence, the implementation of vector competence surveillance programs should be mandatory for different geographic areas, even in the same country.

A CPE occurred at later times in cells inoculated with salivary glands of *Cx. tarsalis* (4 dpi), suggesting that it may not be a competent vector. Alternatively, is cannot be discarded that the salivary glands in this pool had been recently infected, because the CPE in the rest of the body was observed at 1 dpi.

In those cases where ZIKV was found in salivary glands, a CPE was observed at a similar dpi in 5 wild-caught mosquito species, which are therefore potential vectors of ZIKV. Nevertheless, further studies of a possible vector competence barrier to ZIKV in all mosquitoes reported herein are needed, since many factors could be involved in the transmission of the virus as has been suggested^15^.

Furthermore, we found ZIKV in a male pool of *Ae. aegypti*, supporting previous reports in mosquitoes from Brazil and in experimental infections^16,17^; in addition, we also found a ZIKV-positive male pool of *Cx. quinquefasciatus*, what suggests vertical transmission and causes further concern. If male mosquitoes are infected vertically, females from the same mother are probably also infected. Therefore, the number of mosquitoes with the potential to transmit the virus increases: these possibilities can be addressed with further study of their saliva. Also, the finding of three infected *Culex* species could be a major concern and potential complication of vector control programs, because all these species have different breeding sites, can maintain viral populations during interepidemic periods, such as the dry season, and hibernate during colder temperatures.

In conclusion, additional studies of female mosquito saliva from the different species reported in this work are needed to confirm the presence of ZIKV and determine if they have a vector competence barrier to the virus.

## Methods

### Mosquito collection

The collection was performed by mechanical aspiration using an InsectaZooka No. 2888A (BioQuip Products, Rancho Dominguez, CA, USA) inside residences. Mosquitoes were transported to the Entomological Research Unit of Jalisco State Public Health Department. Neighborhoods and blocks were selected based on the reports of the Vector Borne and Zoonotic Diseases Department of Jalisco State on ZIKV human confirmed or probable cases in the area.

Mosquitoes were separated in the laboratory by species and sex into pools of maximum 25 insects. Male pools were frozen at -20°C in 1.5 ml conical tubes containing 250 μL of viral transport medium (phosphate-buffered saline, pH 7.4, containing 30% fetal bovine serum and 2% of penicillin, streptomycin, and amphotericin B). Female mosquitoes from each species-specific pool were dissected under a stereomicroscope. Their body parts (salivary glands, midguts, and rest of the bodies) were distributed into individual pools containing viral transport medium. Some mosquito females were frozen without dissections and were only processed for virus isolation.

### Virus Isolation

Mosquito pools were ground and the resultant homogenates were centrifuged at 10,000xg for 10 minutes. Next, 25 μL of each supernatant was placed into a single well of a 24-welled plate containing *Aedes albopictus* cells C6/36 (ATCC® CRL-1660^™^) or Vero cells (ATCC® CCL-81^™^). The later cell line was used to confirm if isolates were viruses that infect humans. After the inoculum was absorbed for 1 hour at 28°C or 37°C, maintenance medium was added. Cultures were maintained in an incubator at 28°C or 37°C and examined daily for evidence of viral cytopathic effect (CPE) for 5 days. If no CPE was observed, the culture was frozen-thawed once and re-inoculated in a blind passage in a fresh plate for another 5 days. If no CPE was still observed, cultures were discarded.

### Virus Identification

Viral RNA was extracted from the cultures that showed CPE after a single C6/36 passage using a QiAmp Viral RNA Mini Kit (Qiagen, Hilden, Germany). Real time reverse transcription–polymerase chain reaction assays (RT-qPCRs) were carried out in a Light Cycler 480 II PCR platform (Roche Diagnostics, Penzberg, Germany) using Verso 1-step RT-qPCR Kit (Thermo Fisher^™^, MA, USA).

Since in the same area ZIKV, dengue, and chikungunya viruses have been co-circulating, the presence of ZIKV was determined, first, using the primer pair and probe previously reported by Lanciotti *et al.* that can detect 25 genomic copies of the virus^18^. As a positive control of the reaction, we used RNA extracted from a ZIKV strain donated by A.A. Sall (Institut Pasteur at Dakar, Senegal). If the cell cultures showing a CPE resulted negative to ZIKV, then RT-qPCRs for chikungunya and dengue was performed.

### Analysis

We estimated the infection rates (IR) per 1,000 mosquitoes, with the bias corrected by maximum likelihood estimator (MLE) with a skewness-corrected score confidence interval, using the program PooledInfRate v.4.0^19^.

## Footnotes

3 Members of the Mexican Network for Virology who contributed with planning and discussion of data: Carlos F. Arias, Susana López, Rosa María del Ángel, and Victoria Pando-Robles

## Acknowledgments

This work was partially supported by the National Council for Science and Technology (CONACyT)-Mexico. We are in debt with Drs. José Narro Robles, Pablo Kuri Morales, and Cuitláhuac Ruiz Matus, from the Secretaría de Salud of México, for their help and support to carry out this work. We also thank the personnel of Unidad de Investigación Entomologica de Occidente (the Entomological Research Unit of Jalisco State Public Health Department) for their assistance in the collection of mosquitoes and separation of species.

